# Dual-graph deconvolution of enzyme fitness: an interpretable framework to identify and explain catalytic modulators

**DOI:** 10.64898/2026.10.15.759402

**Authors:** Sergio Garcia Busto, Ethan Eschbach, Thomas A Hopf, Chris Sander, Sergio Bacallado, Debora S Marks

## Abstract

Models trained on protein sequences can predict mutational effects, but their outputs represent a convolution of different phenotypes. Isolating stability-independent activity effects is essential for rational enzyme design, allosteric drug design and functional annotation, yet existing pLM-based deconvolution methods lack interpretability. Here, we introduce a hybrid framework that pairs stability predictors with probabilistic graphical models of coevolution. Benchmarked on all enzymes with available multidimensional deep mutational scanning data, our framework isolates stability-independent activity effects with a fidelity that matches analogous pLM-based approaches. Crucially, we reveal biophysically plausible communication pathways that align with previous knowledge and that we validate with experimental data across a diverse set of targets. This hybrid approach merges statistical and geometric deep learning into a zero-shot, explainable tool for targeted engineering and mechanistic discovery.

## 1 Introduction

Enzymes catalyze biochemical reactions with exceptional specificity and efficiency, rendering them indispensable tools across biotechnology and medicine^1,2^. While engineering enzymes with enhanced properties offers immense promise, identifying the precise sites governing catalytic function and explaining how they alter reaction outcomes continue to bottleneck biotechnology^3^. Establishing this mechanistic basis is essential for biological discovery, de-risking engineering campaigns through hypothesis-driven library diversification, and turning model predictions into targeted, high-confidence sequence modifications^4,5^.

Recent advances in machine learning have led the field closer to this goal. For instance, probabilistic evolutionary models trained on deep sequence alignments predict mutation effects with remarkable accuracy, frequently matching or exceeding experimental assays and models trained on the entire protein universe^6–10^. This predictive success stems from the fact that evolutionary sequence conservation natively encapsulates the selective pressures acting on a protein. Nonetheless, this global evolutionary signal represents a complex convolution of overlapping phenotypes, mixing thermodynamic stability, solubility, catalytic activity and regulation into a single score^8^. For instance, while a mutation can be flagged as highly deleterious because it violates a localized functional interaction, standard evolutionary models view this through the same lens as a mutation that forces the entire tertiary structure to collapse^9^.

Resolving this phenotypic mixture is not only important for uncovering the molecular mechanisms driving catalysis, but also for overcoming current limitations in biochemical and biomolecular design^11–15^. For rational enzyme engineering, combining activity-increasing mutations that marginally destabilize a protein with compensatory stability-inducing mutations requires a breakdown of the relative contribution of stability and activity for each mutant^11^. In addition, activity-modulating sites that preserve protein integrity are prime targets for mutagenesis in directed evolution experiments^15^. Similarly, for allosteric drug discovery, deconvolving activity and stability components is essential to identify distal, solvent-accessible pockets that could modulate catalytic activity^16^. Ultimately, accounting for stability as a *condition* for catalysis enables the systematic discovery and inspection of all phenomena that affect activity, including inter-domain or intermolecular functional interactions and shifts in conformational equilibria.

Aiming to address this challenge, recent deep learning frameworks attempt to deconvolve fitness effects by examining the inconsistencies between evolutionary scores from protein language models (pLMs) compared to stability predictors or by partitioning latent representations of pLMs^17–22^. However, because these frameworks rely on black-box pLMs, they cannot explain their predictions nor trace the biophysical pathways through which perturbations alter function. Operating within uncharacterized, high-dimensional latent spaces is also arguably unnecessary for single mutants, given that all approaches ultimately rely on independent stability and evolutionary fitness predictors. This lack of interpretability prevents researchers from discovering the physical interactions giving rise to a phenotype, leaving the underlying residue networks—which could otherwise inform biomolecular engineering, mechanistic discovery, and functional annotation—untapped.

To bridge this gap, we introduce a hybrid framework that pairs stability predictors based on graph neural networks (GNNs) with probabilistic graphical models of protein coevolution (Figure 1). Using a diverse set of enzymes with multidimensional deep mutational scanning (DMS) data, we show that our framework can both recover stability-independent mutation effects and trace how information is transmitted through the structure to affect the active site. We generate plausible information transfer paths and validate them using experimental data for various targets to uncover statistically grounded mechanistic hypotheses that align with prior knowledge. Ultimately, this work moves the field beyond blind functional prediction, providing an end-to-end, fully explainable foundation for targeted enzyme engineering, allosteric drug design, and mechanistic discovery.

**Figure 1:**
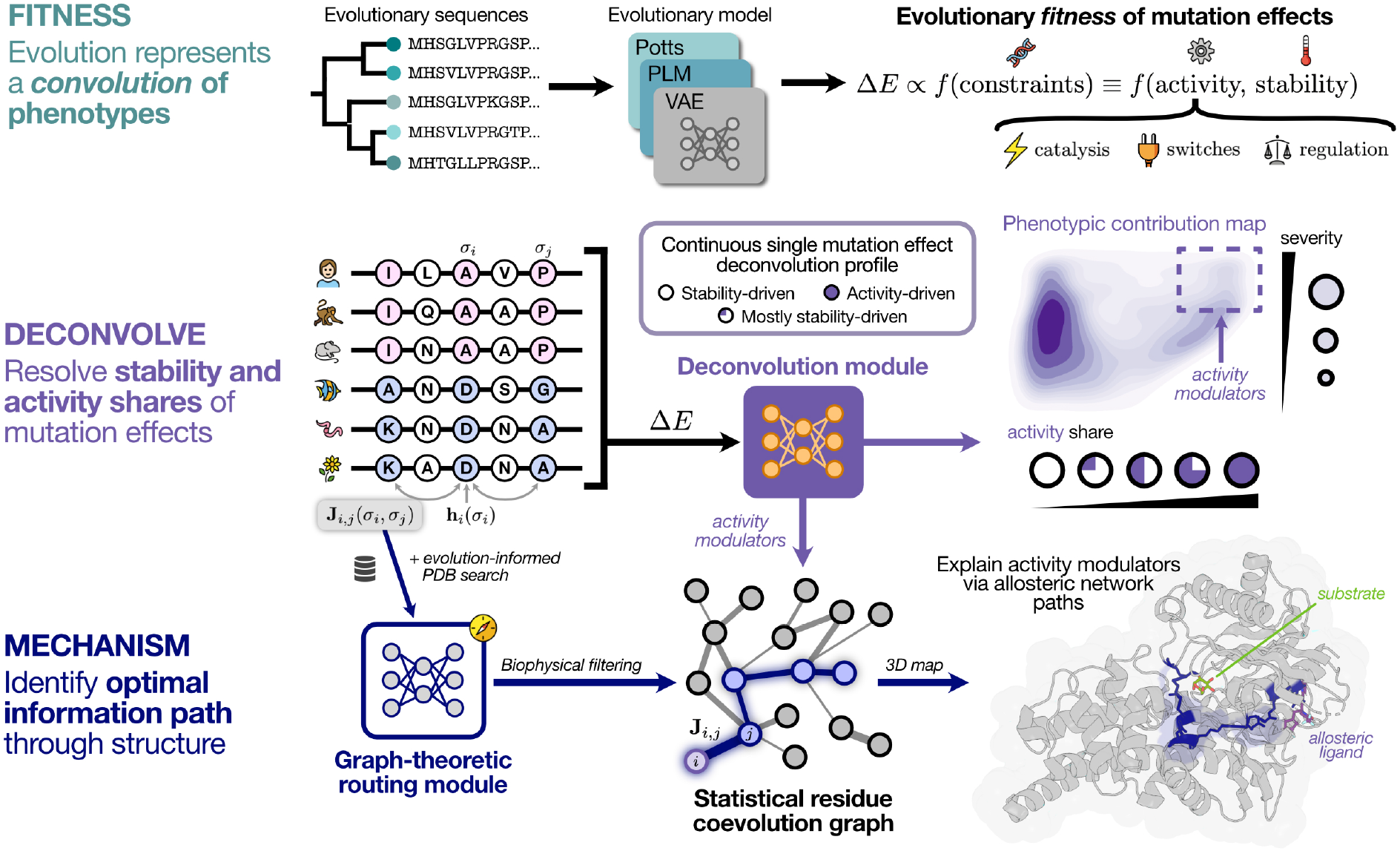
Overview of the proposed dual-graph deconvolution framework. A model of statistical coevolution^23^ serves to predict mutation effects and build a coevolution graph. GNN-based tools are used to identify stability-independent modulators of enzyme activity, which are used as an “information source” to find the most supported information transfer path in 3D space. The outputs of the deconvolution step can be transformed into a phenotypic contribution map for variant annotation (see Methods 4.5). Both the phenotypic contribution map and the information transfer path shown represent a real example of human glucokinase (UniProt ID: HXK4_HUMAN; PDB ID: 3vev). GNN, Graph Neural Network.

## 2 Results

### 2.1 An interpretable latent-variable framework reveals stability-independent catalytic effects

To contextualize the extent to which evolutionary models can deconvolve phenotypes, we first evaluated the predictive performance of leading sequence-based and structure-sequence models across the set of enzymes that are the focus of this study. These represent, to our knowledge, the only enzymes with DMS data for both protein stability as abundance and catalytic activity in the peer-reviewed literature^16,24–31^ (Supp. Table 1; see Methods 4.1). We observed no consistent, statistically significant difference in predicting activity between structure-informed “hybrid” pLMs and alignment-based alternatives like EVE^8^ or EVmutation^6^ (Fig. 2a, b). These observations support massive community benchmarking efforts demonstrating that models leveraging multiple sequence alignments (MSAs) match or exceed the performance of more complex architectures and larger training datasets for predicting activity^10^ and other phenotypes^32^. Interestingly, performance appeared to follow experimental activity-stability correlations, reinforcing the well-documented stability bias of evolutionary models^33,34^.

**Figure 2:**
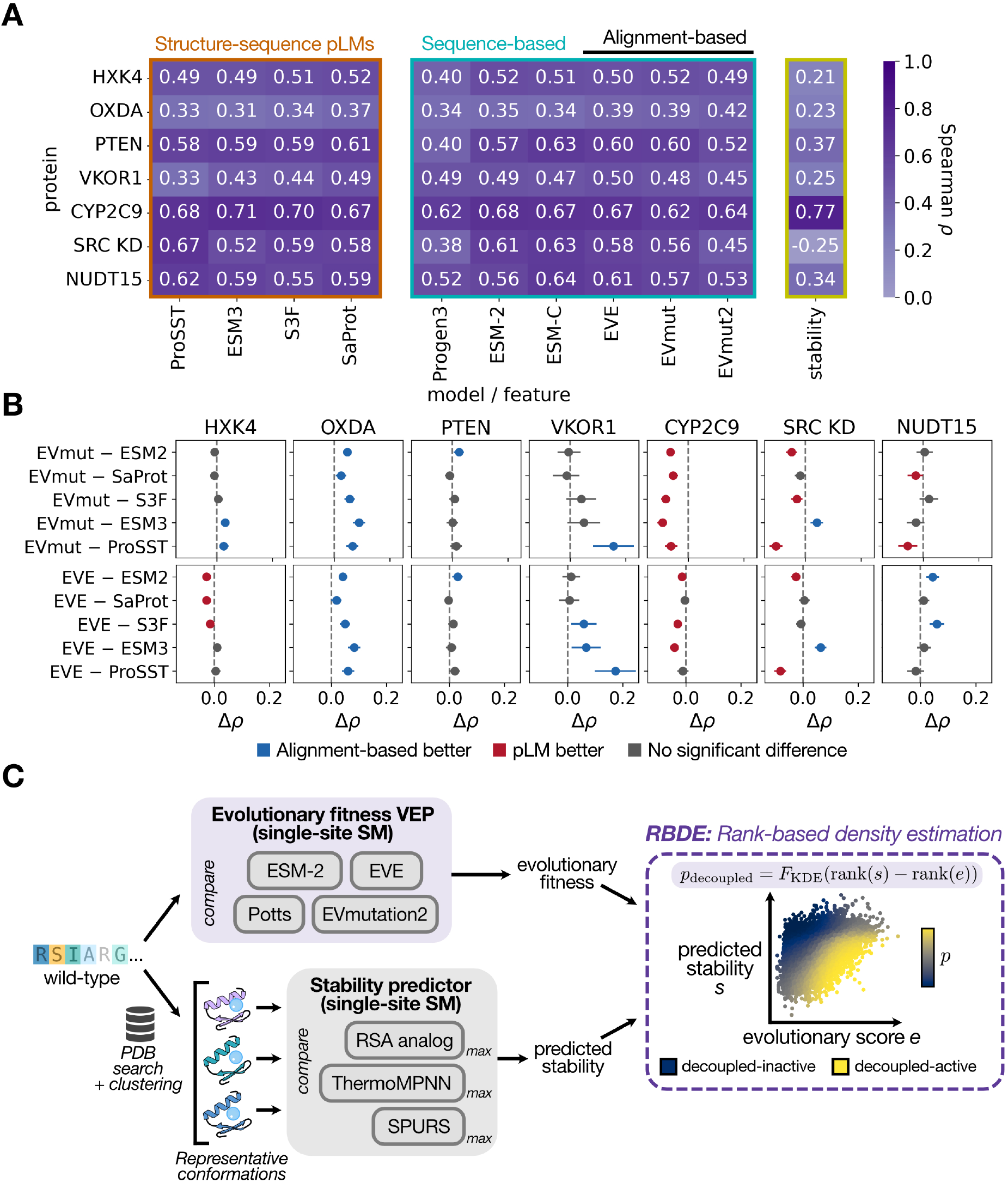
Overview of validation datasets and deconvolution module. (A) Spearman correlations of evolutionary model scores and raw DMS activity data, as well as activity-stability correlations from the multidimensional DMS. Models are classified by structure-sequence pLMs that do not accept an MSA as input, sequence-based pLMs and alignment-based models. Model scores were obtained from ProteinGym^10^. (B) Head-to-head comparison of EVmutation^6^ (top) and EVE^8^ (bottom) with top pLMs. The difference in Spearman correlation coefficient (*ρ*) is reported with 95% confidence intervals, which were obtained by non-parametric bootstrapping of 1,000 samples. Evidence of superior predictive performance of alignment-based models is shown in blue, superior performance of the pLM is shown in red, and no significant difference is colored gray. (C) An overview of the deconvolution module, which evaluates the consistency of evolutionary scores and stability predictors non-parametrically (see Methods 4.3). The predicted stability landscape is built in a conformation-aware manner by recovering homologous structures in the PDB for the target, clustering based on RMSD, and selecting representative conformations. The most destabilizing prediction is kept for each mutant. DMS, Deep Mutational Scanning; pLM, protein Language Model; MSA, Multiple Sequence Alignment; PDB, Protein Data Bank; RMSD, Root Mean Squared Deviation; SM, saturation mutagenesis.

#### 2.1.1 Statistical models of protein coevolution recover stable-but-inactive variants similarly to protein language model-based alternatives

In order to identify stability-independent mutation effects, we built a non-parametric, conformation-aware structure-sequence module that combines (i) a stability predictor and (ii) an evolutionary fitness predictor, to produce a score for a mutant protein being stable-but-inactive (Fig. 2c, see Methods 4.3). For evolutionary models, since hybrid pLMs and sequence-based models have comparable predictive power (Fig. 2a, b), we compared the ESM2 pLM of different parameter sizes to: a statistical model of protein coevolution, EVmutation^23^; an alignment-based VAE, EVE^8^; and an alignment-based autoregressive model, EVmutation2^35^. For stability predictors, we compared ThermoMPNN^36^ and SPURS^37^, two state-of-the-art GNN-based models used in prior functional deconvolution efforts^19,21^. Given the power of residue buriedness metrics to predict variant effects^38–40^, we also included a relative solvent accessibility (RSA)-based baseline that incorporates mutation-specific penalties (see Methods 4.2). Interestingly, this baseline predicted stability similarly to the more complex counterparts for most targets (Supp. Fig. 1).

Baseline evolutionary models were consistently outperformed by “joint” models, both in their ability to recover stable-but-inactive variants by NDCG, AUPRC and AUROC, and by global correlation to a model only built with experimental data (Fig. 3a, b; Supp. Fig. 2). Crucially, we observed no statistically significant difference between ESM2 and EVmutation in recovering stable-but-inactive variants by global metrics, and no consistent difference by top-heavy metrics like NDCG (Fig. 3c). We found SPURS to be marginally superior to ThermoMPNN across enzymes, surpassing the RSA analog baseline (Fig. 3d, Supp. Fig. 3). We additionally investigated the ability of our approach to isolate mutants that increase activity at the expense of stability, which the deconvolution module natively predicts via the complement of the deconvolved probability (Fig. 2c, see Methods 4.3). Intriguingly, we observed no consistent improvement of joint models over evolutionary models for this task, suggesting an increased difficulty of predicting stability for mutants that are not heavily penalized by the evolutionary model (Supp. Fig. 4).

**Figure 3:**
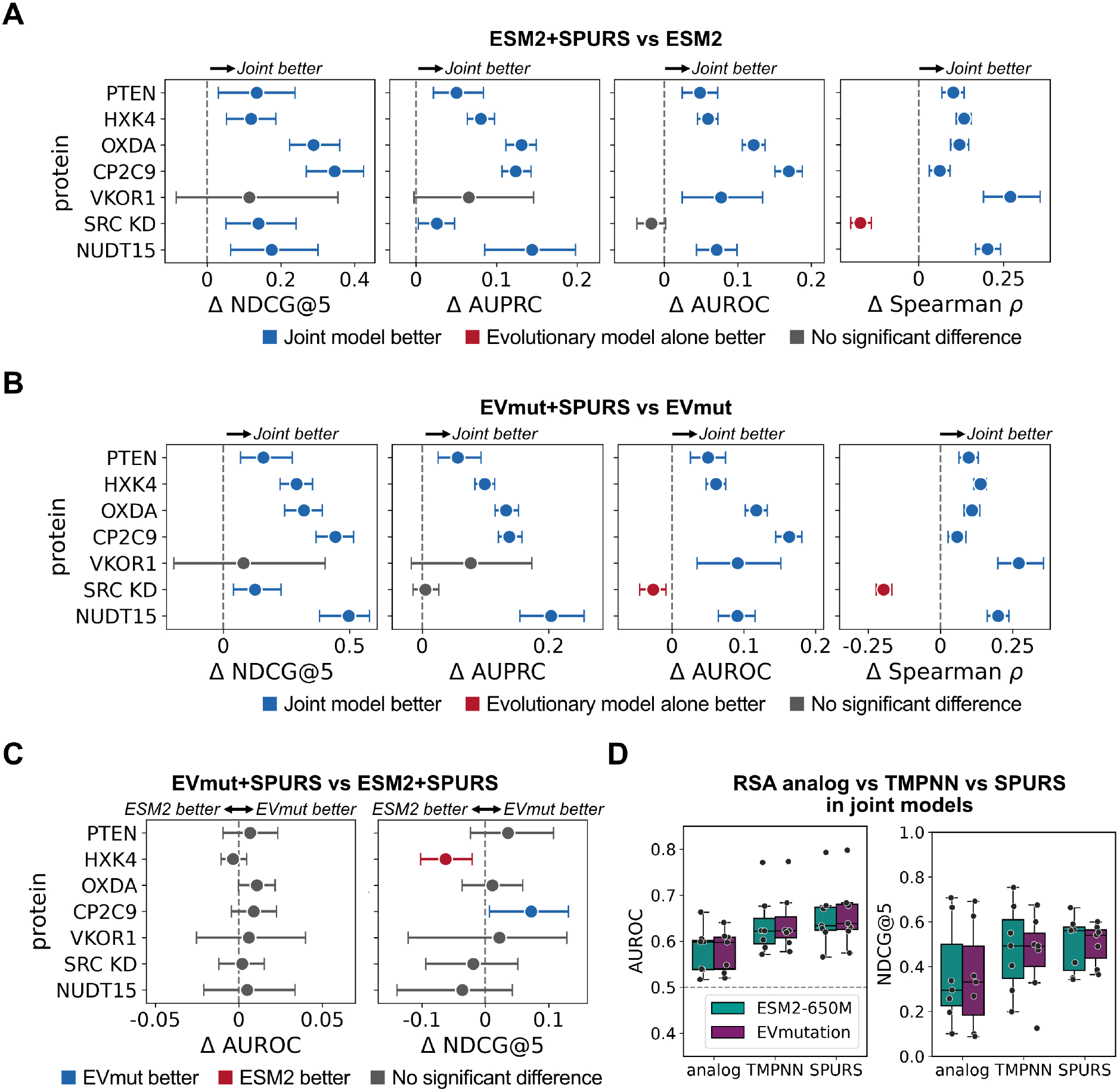
Stable-but-inactive variant recovery performance across model combinations. (A) Head-to-head comparison between the joint model composed of ESM2-650M and SPURS as the evolutionary fitness and stability predictors, respectively, and ESM2 alone. (B) An analogous comparison to (A) with EVmutation as the evolutionary fitness predictor, highlighting that joint models offer a significant advantage over evolutionary models alone. (C) Joint model comparison between EVmutation- and ESM2-SPURS models by global (AUROC) and top-heavy (NDCG@5) metrics, suggesting EVmutation and ESM2 recover stability-independent mutation effects comparably. (D) Joint model comparison between stability predictors across ESM2 and EVmutation, suggesting a marginal improvement when using SPURS (see Supp. Fig. 3). For (A), (B) and (C), the difference in each metric is reported with 95% confidence intervals, which were obtained by grouped non-parametric bootstrapping of 1,000 samples. AUROC, Area Under the Receiver Operating Characteristic curve; AUPRC, Area Under the Precision Recall Curve; NDCG, Normalized Discounted Cumulative Gain.

Beyond isolating variants with extreme deconvolution scores, the deconvolution module can also be applied to the task of clinical variant annotation. To enable this, we derived a parameter-free 2D coordinate space that maps any substitution along continuous stability and activity axes, establishing a simple and transparent scheme for variant interpretation (Fig. 1; see Methods 4.5).

#### 2.1.2 Deconvolved scores isolate evolutionarily conserved functional sites

If the deconvolution module successfully decouples activity from stability, predicted stable-but-inactive variants should, at least partly, map to documented functional sites. While previous studies used a single reference structure for various analyses^17,19,20^, this overlooks key aspects of protein function, including the dynamic nature of catalysis represented by alternative conformations, the information obtained from substrate analogs and intermolecular interactions, and the complementary evidence from homologous enzyme structures. To address these gaps and consider the full functional profile of the targets, our framework maps all functional interactions— including protein-protein interactions and relevant small molecule-binding pockets—leveraging all structures from homologous enzymes (see Methods 4.4).

As reported in previous work^20^, we observed a clear enrichment in functional sites across targets (Fig. 4a). Importantly, while this is foreseeable when using standard evolutionary models^17,41^, the deconvolved signal isolated sites that retained wild-type-like stability levels when mutated (Fig. 3b). Additionally, we observed comparable recovery performance between ESM2 and EVmutation at the lowest percentiles of joint model scores, arguably the most important data regime for discovery and engineering campaigns (see Section 2.2). We found notable diversity in the nature of functional interactions, including protein-protein interactions (PPIs), allosteric pockets, and active site-binding interactions, owing to differences in evidence for each enzyme in the Protein Data Bank (Fig. 4c). Thus, as expected^19,41^, deconvolved mutant sites were not confined to the active site alone (Fig. 4d).

**Figure 4:**
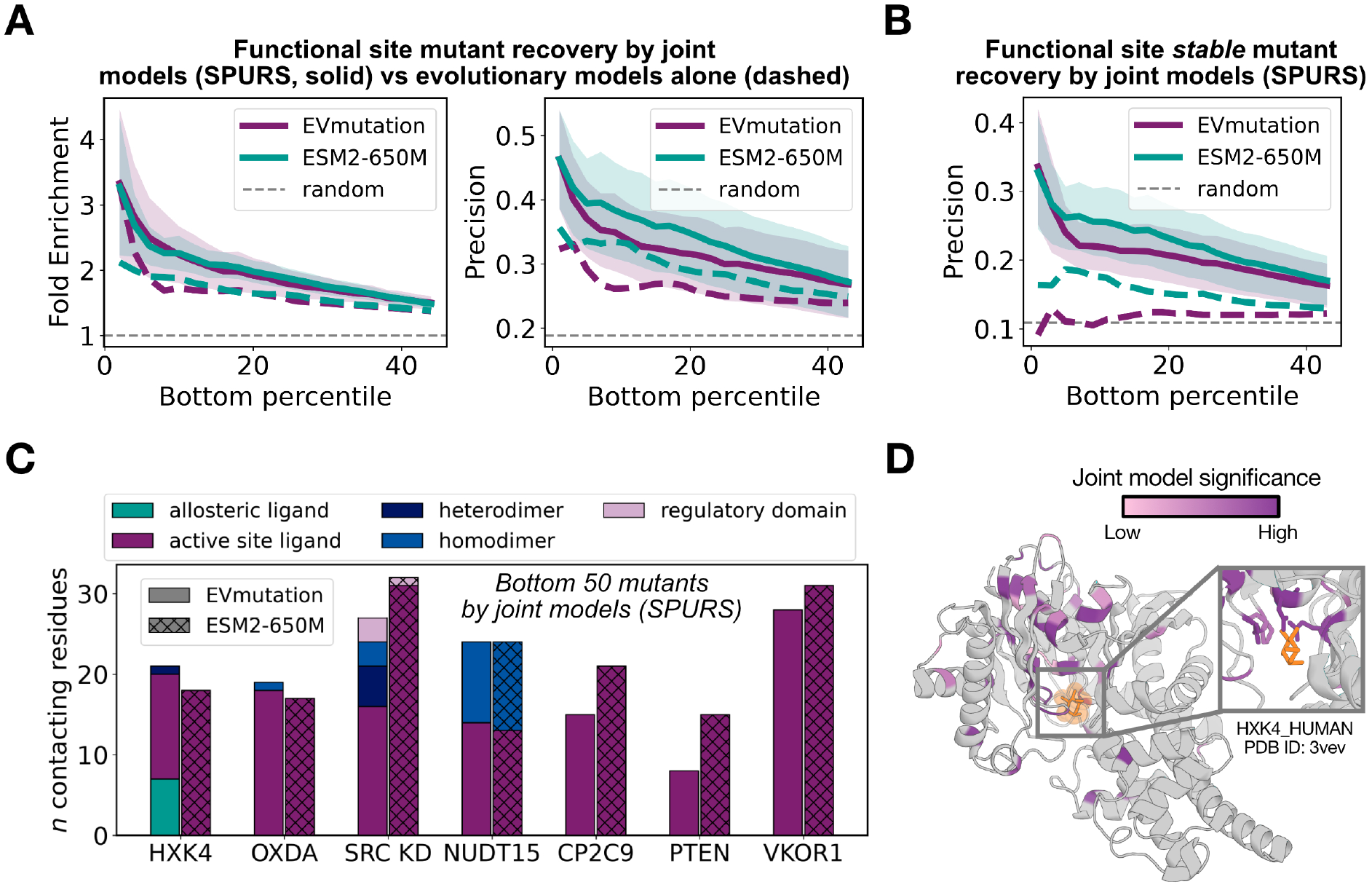
Evolution-informed functional site recovery from the deconvolution module across targets. (A) Recovery of functional sites as mean fold enrichment (left) and precision (right) when screening for stable-but-inactive scores (see Fig. 2c) using SPURS as the stability predictor and either EVmutation or ESM2 as the evolutionary model. Performance from evolutionary models alone is shown by dashed lines of corresponding colors. (B) An analogous plot to (A) assessing the recovery of variants within functional sites that specifically retain wild-type-like stability, where joint models are specifically efficient. (C) Functional site characterization within the 50 most significant mutants of joint model scores. (D) Unique sites corresponding to the 40 most significant deconvolved mutants of the EVmutation-ThermoMPNN model highlighted within a glucokinase structure, illustrating that identified sites are not confined to the active site alone. Shaded regions in (A) and (B) represent the standard error of the mean.

### 2.2 Phenotype deconvolution and graph-theoretic routing generate plausible mechanistic hypotheses of enzyme allostery

We hypothesized that allosteric or activity-modulating signals propagate along physically contiguous residue networks, whose information transfer capacity could be approximated by pairwise evolutionary couplings^23,42,43^. We therefore formulated this signaling as a graph-theoretic routing problem by projecting coevolutionary coupling strengths onto a 3D residue-residue contact map (Fig. 5a). Anchored by our phenotype deconvolution framework, which identifies the top 1% of stability-independent mutational effects as information sources, this approach isolates the most statistically and physically supported pathways transmitting allosteric signals to the active site (see Methods 4.6).

**Figure 5:**
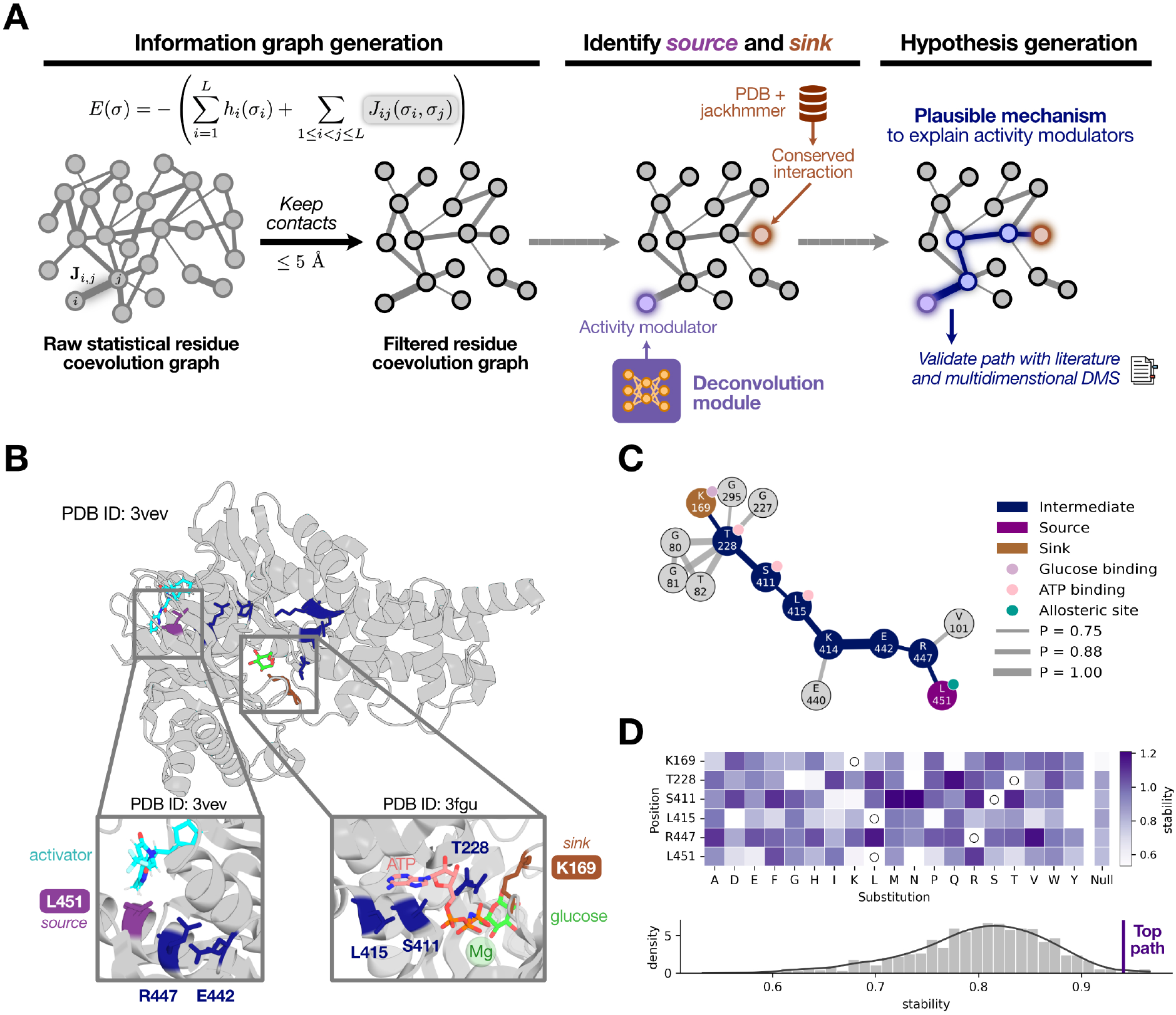
An overview of the network routing module and the case study of glucokinase activation by allosteric ligands. (A) The network routing module starts with a coevolution graph from EVcouplings^23^, which is filtered to keep physically contiguous pairwise relationships. Activity-modulating sites from the deconvolution module and evolutionarily conserved active site residues form the information sources and sinks, respectively. Finally, the top-*k* shortest paths are traced between all potential source-sink pairs, which we validate using experimental data and known mechanistic information from literature (see Methods 4.6). (B) The top path linking a substrate-contacting residue in human glucokinase (HXK4_HUMAN) and an allosteric activator binding site. (C) The filtered coevolutionary graph highlighting the path in (B) (dark blue). (D) *Top*: experimental stability of path constituents compared to the expected stability of a set of residues of similar buriedness (*±*0.1 RSA). *Bottom*: the null distribution of 1,000 residue sets of the same length and similar buriedness, showing a significantly higher stability in constituents of the selected path (*p* = 0.007). Residues K414 and E442 were excluded for visualization, but the entire path also has significantly higher stability than the null (*p* = 0.036, see Methods 4.6). RSA, Relative Solvent Accessibility.

#### 2.2.1 Coevolutionary network routing recapitulates and extends the mechanism of glucokinase activation

Human glucokinase (HXK4 or GCK) harbors a well-characterized allosteric pocket that binds activator ligands to regulate catalytic activity^44,45^. The primary mechanism of GCK activation involves shifting the enzyme’s conformational equilibrium toward its closed, active state via a large-scale domain movement^46^. This macroscopic transition restructures the glucose- and ATP-binding cleft, which enhances glucose affinity to drive catalysis^47^. The phenotype deconvolution module successfully identified this regulatory site via variant L451W—a stable-but-inactive mutation^25^ located at a position shown to be required for locking GCK into its active conformation^46,47^.

Originating from L451 as the information source, the network routing module resolved an 8-residue pathway connecting the allosteric pocket to the catalytic glucose-binding residue K169 with a mean coupling probability of 0.82 (Fig. 5b, c; Supp. Table 2). Crucially, the routed network recapitulated the known mechanism, as it traversed three ATP-contacting residues (228, 411, and 415) alongside residue 442—a critical hinge residue that links the large domain to the *α*13 helix and pivots during domain closure^45,48,49^ (Fig. 5b). In addition, the path included K414, a clinical variant site known to impair affinity for both glucose and ATP^45,50^ and reduce protein abundance^25^. Constituent path positions were enriched for high stability when compared to length- and RSA-matched residue sets (Monte Carlo Randomization, *p* = 0.036; see Methods 4.6), and excluding the structural hinge residue E442 and site K414, over 95% of substitutions retained the wild-type phenotype (*p* = 0.002, Fig. 5d; Supp. Table 3). This profile suggests that alongside global shifts in the conformational equilibrium, GCK may employ a dedicated, mutationally tolerant corridor that directly couples glucose and ATP binding, physically integrated with the allosteric pocket through the helix *α*13 hinge^44,49^. We note that pathway residues were not enriched for lower activity, suggesting a switch-like behavior that is only triggered from the allosteric site^46,47^.

#### 2.2.2 Information transfer pathways originating from protein interfaces exhibit mechanistic heterogeneity and align with known biology

Since regulatory proteins and domains tune catalytic function, their interfaces with target domains serve as natural activity modulators. Network routing across enzymes from our benchmark set with PPIs—GCK–GKRP^51^, OXDA^52^, NUDT15^53^, and the SRC kinase domain^54^—successfully resolved interface-to-active site communication pathways across all evaluated assemblies except the SRC heterodimeric interface (Fig. 6, Supp. Tables 2-4). Intriguingly, functional validation using DMS data revealed two distinct biophysical modes of signaling that align with the known functional requirements of each target.

**Figure 6:**
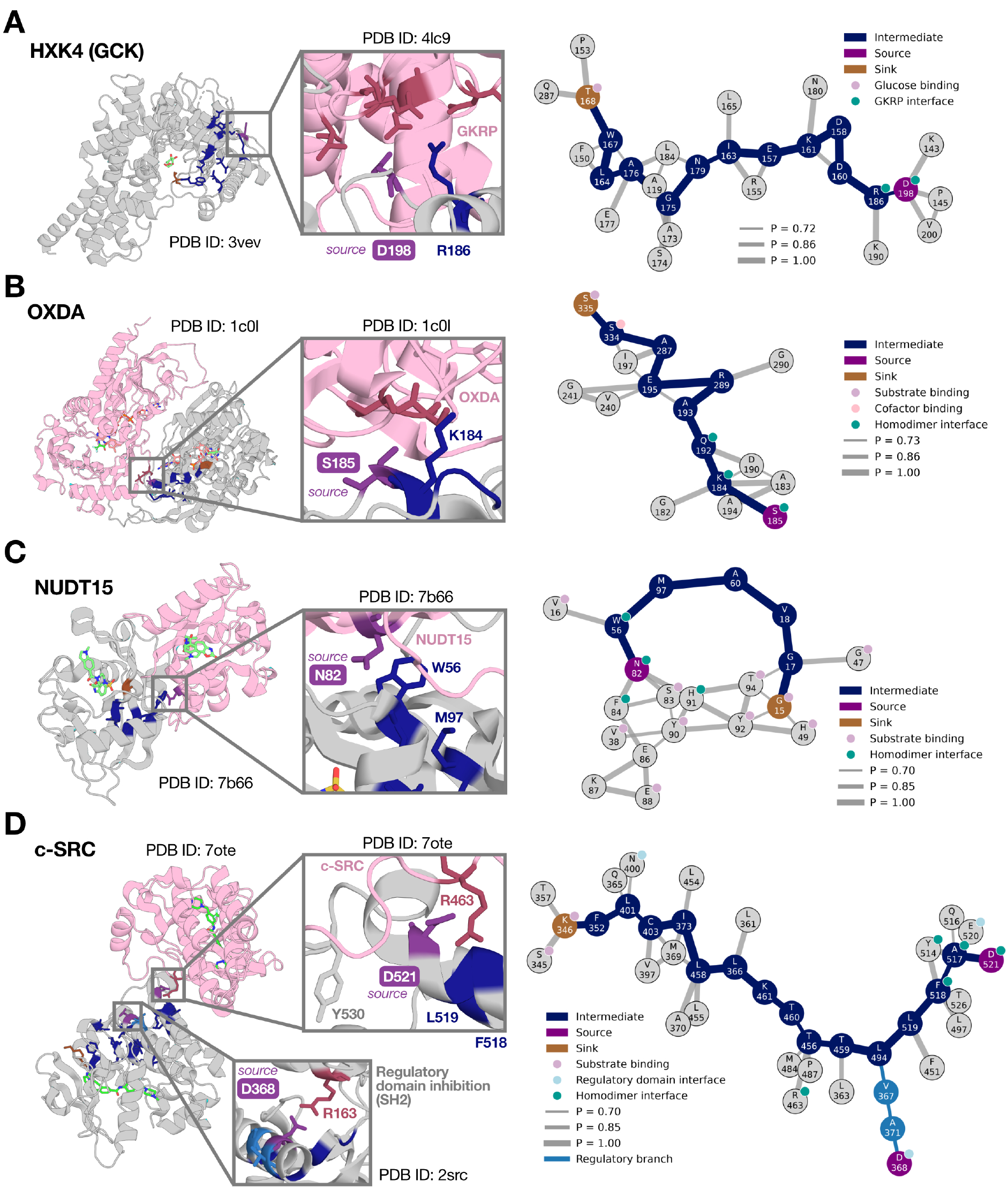
Information transfer paths that can explain how functional protein interfaces affect the active site can be identified *de novo*. (A-D) Structural mapping of selected paths (left) and paths highlighted within corresponding coevolutionary graphs (right, dark blue), for human glucokinase (A), D-amino acid oxidase (B), nudix hydrolase (C) and the kinase domain of Src kinase (D). The latter has two overlapping networks: a primary homodimer interface network (dark blue) and a branching path stemming from the regulatory domain interface (light blue). Both information sources represent inhibitory interactions, but they overlap via a hydrophobic pocket composed of residues L519, L494, T459 and L363, which are thought to mediate signaling^58^. The inhibitory dimerization event mediated by D521 promotes the phosphorylation of Y530 *in trans*, a key event in the signaling lifecycle of Src^54^. Information sources for each highlighted path are colored purple and were identified within the top 1% of deconvolved mutants for each target (see Methods 4.6). Information sinks are colored brown.

In GCK and OXDA, where inter-domain interactions are not essential for baseline enzyme activity^51,55^, we found that regulatory signals travel through thermodynamically shielded conduits. In the GCK–GKRP complex, phenotype deconvolution highlighted interface variant D198W, a mutant that disrupts GKRP binding^51,56^, from which routing resolved a 13-residue pathway to the active site position T168 (mean coupling of 0.94; Fig. 6a). This conduit exhibited broad mutational tolerance, with over 91% of substitutions retaining wild-type-like abundance (*p* = 0.001) and a significant enrichment for stability (*p* = 0.001; Supp. Fig. 6, Supp. Table 3). Similarly, in the non-obligate dimer OXDA^52^, routing yielded a 9-residue conduit (mean coupling of 0.98, Fig. 6b) significantly enriched for high stability (*p* = 0.001; Supp. Table 3) and the wild-type-like stability phenotype (71% vs. 47% baseline proportion, *p* = 0.001). This profile was identifiable by computational predictors (SPURS, *p* = 0.001), confirming that these paths can be discovered *de novo*. Crucially, constituent residues along these paths showed no enrichment for reduced catalytic activity nor evolutionary scores, aligning with the fact that these regulatory signals modulate activity but are not essential for catalysis^51,55^.

Conversely, in NUDT15 and SRC, our framework resolved mutationally sensitive sub-networks where structural integrity and activity are coupled, and which recapitulate their corresponding functional modes. In NUDT15, a strictly obligate dimer^53^, we discovered a dense coevolutionary network characterized by first- and second-degree interactions due to the proximity of the active site and dimer interface (Fig. 6c). Intermediate residues of an isolated 7-residue conduit with a mean coupling probability of 0.99 were significantly enriched for catastrophic variants exhibiting below-wild-type activity and stability (*p* = 0.047; Supp. Table 3), aligning with low evolutionary scores (*p* = 0.03). In SRC, paths traced from inhibitory information sources—the intramolecular SH2 clamp^57^ and the intermolecular enzyme–substrate dimer interface^54^—converged on a shared 12-residue sub-network (Fig. 6d). Remarkably, despite originating at sites of autoinhibition, these pathways coupled to the active site via the C-lobe hydrophobic pocket (residues L363, T459, L494, and L519), which gates a myristate-dependent dimerization event critical for Src signaling that has eluded structural determination due to its lipidated, disordered nature^58^.

This sub-network was enriched for catastrophic mutations compared to RSA-matched null sets (*>* 56% of variants, *p* = 0.002, a 65% increase over baseline; Supp. Table 3), suggesting an essential allosteric conduit. This high mutational sensitivity is consistent with the presence of residues from helix *α*F (T456, L458 and T459), which anchors the functionally critical catalytic (C) and regulatory (R) spines^16^.

#### 2.2.3 *De novo* prediction of a solvent-accessible catalytic dial in a human phosphatase

We next evaluated the ability of our framework to discover potential catalytic switches *de novo* in enzymes lacking structural evidence of regulatory sites. Focusing on human phosphatase and tensin homolog (PTEN), we anchored the routing module’s information sink to residues contacting the co-crystallized transition-state analog vanadate^59^. Ranking the candidate information sources by relative solvent accessibility (RSA) prioritized residue D77 within the catalytic phosphatase domain. The deconvolution module predicted a strong activity-modulating phenotype at D77, as over half of all substitutions scored in the top decile, with three variants falling within the top percentile. One of the latter variants, D77W, maintained wild-type activity despite reduced stability, potentially functioning as a “catalytic dial” that increases enzymatic activity per unit of protein^29^.

Tracing the shortest pathway from D77 to the active site resolved a 6-residue corridor (mean coupling score of 0.82) traversing the core of the catalytic domain, with all four intermediate residues deeply buried (RSA *<* 0.04, Fig. 7a, b). Pathway residues were enriched for protein stability by DMS (*p* = 0.005, Fig. 7c) and had a significantly higher proportion of variants of wild-type-like stability compared to RSA-matched background sets (*>* 75%, *p* = 0.004). Our results suggest that this switch operates via a biophysically shielded corridor similar to those observed in GCK and OXDA. Furthermore, intermediate residues displayed near-significant enrichment for computationally predicted stability by SPURS (*p* = 0.065), highlighting that functional activity dials can be identified *de novo*. Interestingly, the path was also significantly enriched for stable-but-inactive variants (*p* = 0.013), enforcing the mode of action of this subnetwork. Together, these results demonstrate that our framework enables the identification of activity-modulating sites targetable by biologics or small molecules, while generating biologically plausible mechanistic hypotheses.

**Figure 7:**
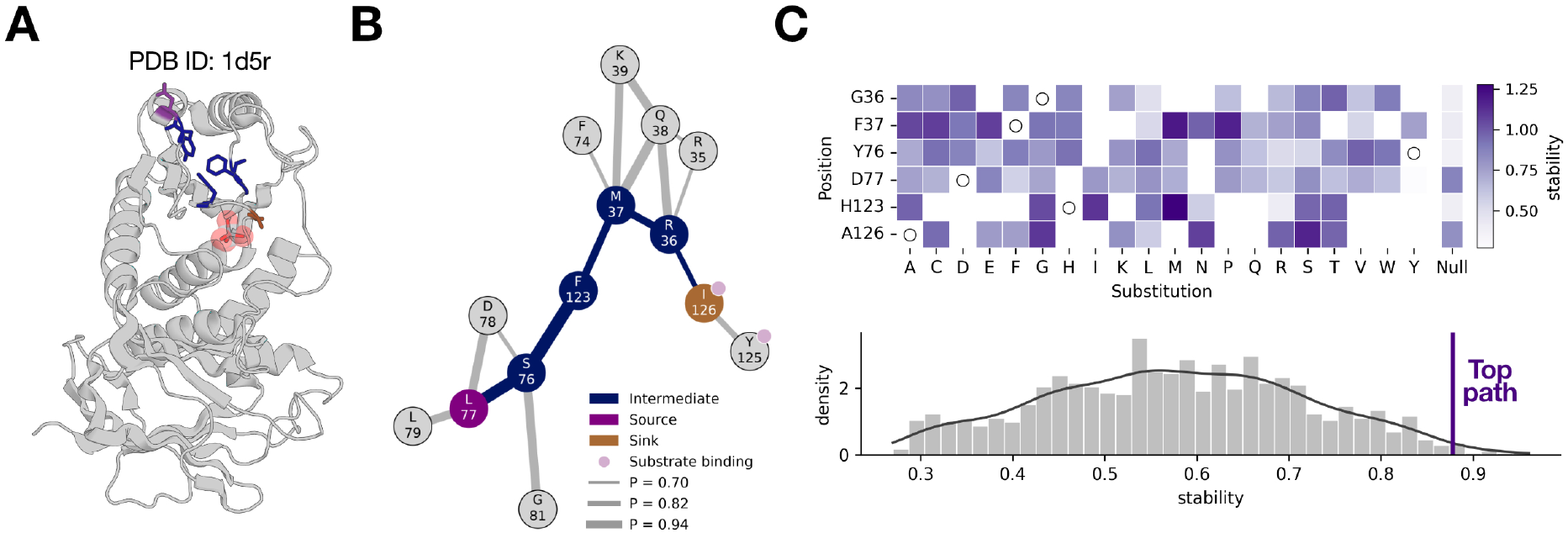
Discovering a catalytic modulator and its mechanism *de novo* in human phosphatase and tensin homolog (PTEN). (A) The top path linking the most solvent-accessible predicted activity-modulating site (D77) and a substrate-binding residue. (B) The corresponding information transfer path with the top path highlighted in dark blue. (C) *Top*: the stability profile of path constitutents compared to the stability expected from a residue of similar buriedness (*±* 0.1 RSA). *Bottom*: a null distribution of 1,000 residue sets of the same length and buriedness, showing a significantly higher stability of the top path (*p* = 0.005, see Methods 4.6). RSA, Relative Solvent Accessibility.

## 3 Discussion

Although pLMs currently dominate variant effect prediction, explicit probabilistic graphical models perform comparably while offering mathematical interpretability^10,23,32,60^. Here, we demonstrate that pairing graph neural network-based stability predictors with statistical models of coevolution can (i) deconvolve stability from evolutionary fitness and (ii) generate plausible mechanistic hypotheses of enzyme function and regulation. Our framework uncovered two distinct intramolecular information transfer modes: stability-enriched conduits that can buffer mutation effects, and core activity-stability coupled sub-networks that, when mutated, are likely to abolish catalysis through disrupting protein folding. We generate pathways in various targets and show that intermolecular interactions, regulatory domain interfaces and allosteric ligand-mediated activation can be identified *de novo* and potentially explained, while aligning with known mechanistic information.

The different DMS enrichment patterns observed align with the nature of the information sources and path intermediates. Paths traversing obligate interaction residues show enrichment in the “catastrophic” phenotype, whereas paths originating from regulatory or non-obligate interfaces show a stability enrichment and no activity deficit. We hypothesize that the stability-enriched signature reflects path residues that accommodate the conformational changes that transmit regulatory signals. This manifests as mutational tolerance in stability by DMS, highlighting the ability of the path to maintain protein integrity and effectively buffer perturbations. In the case of the GKRP–GCK path, which connects the interface of the GKRP inhibitory protein to the GCK active site, disrupting an intermediate would be expected to *relieve* inhibition, since it decouples the active site from the inhibitory signal. A falsifiable test of this hypothesis is therefore a conditional experiment: mutating a predicted intermediate in the presence of GKRP should abolish its effect while maintaining baseline GCK activity.

Identifying these sub-networks uncovers actionable principles for enzyme design, allosteric targeting and mechanistic discovery. The phenotypically coupled sub-networks define machinery that should be preserved during library diversification or potentially co-transplanted alongside the active site during *de novo* design^61^. Conversely, stability-enriched conduits contain ideal target sites for mutagenesis, which evolutionary models alone might deprioritize due to high evolutionary conservation. Rather than directly mutating substrate-contacting residues in directed evolution campaigns, targeting residues along these paths offers sites that specifically modulate catalysis, preserving structural integrity and thus enabling a more efficient exploration of the catalytic landscape^3,5^. Furthermore, where these decoupled conduits terminate at surface-accessible sites, they expose allosteric pockets suitable for drug or protein binder design^13,16^. Importantly, our framework can identify these two functional modes *de novo* by evaluating enrichments in predicted stability and evolutionary fitness, demonstrating its end-to-end and interpretable nature.

Intriguingly, many information sources identified by the deconvolution module—namely the GCK-GKRP, SRC and NUDT15 interfaces and the PTEN catalytic switch—were not classified as stable-but-inactive by DMS. This might stem from the fact that cellular abundance is an indirect measure of protein stability given its sensitivity to protein degradation pathways. Consequently, relying on abundance-based ground truth labels could artificially underestimate the performance of phenotype deconvolution methods^20,21^. Alternatively, this could suggest that solvent-accessible, activity-modulating sites do not necessarily manifest a stable-but-inactive phenotype upon mutation, potentially allowing our framework to identify sites that DMS experiments miss. Our results also suggest that the routing module alone could reveal functional paths independently of the deconvolution step. For example, in SRC, the residue N400 could trace a shorter sub-network linking the regulatory domain interface and a substrate-binding residue, yet was excluded because no single mutation at that site passed the top 1% deconvolution threshold (Fig. 6d). Similarly, R463 was not highlighted as a potential information source, but forms a key interaction with D521^54^ and is indirectly coupled to key functional sites (see Section 2.2.2).

Despite promising results across our enzyme validation set, our approach has several limitations. First, inferred pathways represent biophysical hypothesis generators rather than mechanistic proofs, and may act together with other residues, paths or structural mechanisms such as modifying the conformational equilibrium. Therefore, validating specific conduits requires complementary evidence. Second, because dual DMS data only exists for a small set of enzymes, determining whether these mechanisms are prevalent across targets requires broader multi-phenotype experiments. Third, although the underlying principle of our deconvolution module aligns with existing methods^18,19^, evaluating the diagnostic capacity of our two-dimensional scale requires benchmarking against clinical databases. Lastly, using the Protein Data Bank to confirm whether an identified functional pocket represents a protein interface is limited by available experimental structures. Integrating computational interface predictions^62^ or alignments^63^ could extend this validation beyond solved complexes, revealing functional surface sites that would otherwise be missed. Overall, our framework unifies statistical and geometric deep learning and offers an explainable roadmap for automated catalyst engineering and therapeutic design.

## 4 Methods

### 4.1 Data processing

Activity and stability values for the enzymes within the validation set (UniProt IDs: HXK4_HUMAN, CP2C9_HUMAN, OXDA_RHOTO, PTEN_HUMAN, VKOR1_HUMAN, SRC_HUMAN, NUD15_HUMAN) were obtained from the original sources, as well as the wild-type ranges^16,24–31^. Where reasonable minimum wild-type experimental values were unavailable, the threshold was computed using a Gaussian mixture approach as reported previously^21^. Specific activity (see Section 2.1, Supp. Fig. 4) was defined as

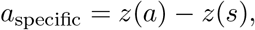

where *z* represents median absolute deviation (MAD)-based standardization, activity *a* and stability *s*. The top 35% of specific activity mutants were taken amongst mutants with at least wild-type-like activity, ensuring that the average proportion of high specific activity variants is sufficient to measure recovery (>15%). The number of stable-but-inactive and high specific activity mutants are shown in Supp. Table 1.

Structures for each target were obtained in a conformation-aware manner (see Fig. 2c) using the Protein Data Bank (PDB) with records up to 2025-08-06. Structures were selected by (i) collecting all structures annotated with the target’s UniProt ID (ii) conserving structures with *>* 80% sequence coverage, and (iii) clustering based on Root Mean Squared Deviation (RMSD) using average-linkage hierarchical clustering and a cutoff of 6 Å. Within the remaining clusters, the structure most cited in existing literature was chosen as a cluster representative. While the Src kinase domain (PDB IDs: 2src, 1y57) and HXK4 (PDB IDs: 3vev, 4dch, 1v4t) had multiple structures, a single structure was used for all other targets (CP2C9_HUMAN, 1og2; OXDA_RHOTO, 1c0p; PTEN_HUMAN, 1d5r; VKOR1_HUMAN, 6wv3; NUD15_HUMAN, 5lpg).

### 4.2 Evolutionary and stability model scores

Alignments were constructed following EVmutation guidelines, and default configurations were used^6,8,35^. ESM-2 model scores were obtained through positional log-likelihoods^64^ using evedesign^35^. ThermoMPNN^36^ and SPURS^37^ were also run with default configurations using the provided Colab notebook and repository, respectively. The sign of ΔΔ*G* predictions was reversed to directly represent stability. Where multiple structures are considered, the most stable prediction was taken. Given that the deconvolution module is nonparametric and relies on rank-based density estimation, no additional transformation was required (see Methods 4.3).

The Relative Solvent Accessibility (RSA)-based analog was built using FreeSASA^65^. RSA was obtained by dividing Solvent-Accessible Surface Area (SASA) by the empirical residue maximum SASA^66^. by computing dissimilarity scores *d* for each single mutant *u*,

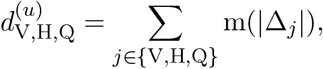

where m is min-max scaling, V is volume^67^, H is hydrophobicity (Eisenberg-Weiss scale^68^) and Q is charge at pH 7.4. Scaling was performed across all 20 *×* 20 substitutions. Then a buriedness score *b* was calculated for each position *i* using the minimum RSA across conformations:

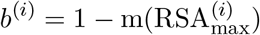

The final stability analog *g* for substitution *u* at position *i* was calculated by

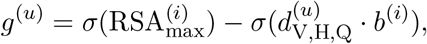

where *σ* represents standard scaling. For membrane-spanning proteins, an additional penalty *l* was added to consider the lipid environment of each residue and account for electrostatic and hydrophobicity alterations in the membrane. First, the structure was oriented using the PPM 3.0 server^69^ to estimate the location of the membrane. The resulting structure was used to split SASA estimates into water, lipid interface and bilayer core environments (32 Å thickness, interface *±*1.9 Å). The penalty can be expressed as

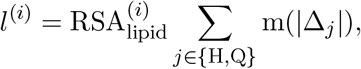

where

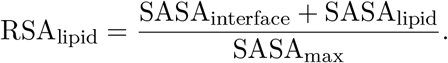

Upon multiple conformations, the minimum of RSA_lipid_ is also taken. Therefore, the final stability analog for membrane proteins was calculated as

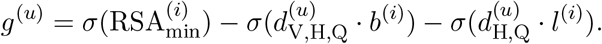

### 4.3 Deconvolution module

The deconvolution module evaluates the consistency of predicted stability and evolutionary fitness (Fig. 2c). For the variants scored by both models, each score was converted to a fractional rank by average ranking and division by the number of variants with both scores, giving *r* ∈ (0, 1].

Therefore, for mutant *u*, the rank difference *v* is simply

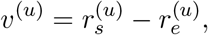

where *s* is the stability prediction and *e* is evolutionary fitness. Thus, positive *v* implies the stability rank exceeds the evolutionary rank amongst all evaluated single substitutions. A Gaussian kernel density estimate 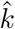, with bandwidth set by Scott’s rule, was fit to the rank differences at or below the upper Tukey fence, *v* ≤ *Q*_3_ + 1.5 × IQR. The lower tail was retained, so the extreme positive differences did not determine the bulk density. Let *F* be the cumulative distribution function of this fitted density, obtained by numerical integration of 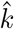. *F* was evaluated for every variant, including those excluded from the fit, and the deconvolution score is

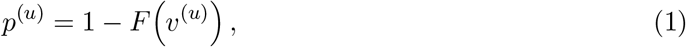

with *p* clipped to [0, 1]. Thus *p* is the survival function of the bulk model: variants whose ranks agree across predictors fall near the center of 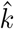 and receive an intermediate score (*p* → 0.5); variants with a higher stability rank than evolutionary rank fall in the right tail and are scored as stable-but-inactive (*p* → 0); variants with a higher evolutionary rank than stability rank fall in the left tail and are scored as high specific activity (*p* → 1). Recovery of high-specific-activity variants was assessed by screening high scores. The same procedure was repeated with experimental activity as *e* and experimental stability as *s*, giving an experimental deconvolution score for each variant. These scores were then correlated with the model-derived deconvolution scores across the full variant set (Spearman Correlation Coefficients, Fig. 3a, b).

### 4.4 Extracting evolutionarily conserved functional contacts from the Protein Data Bank

Query sequences were searched with JackHMMER^70^ (HMMER 3.4) against the SIFTS PDB chain set (records until 2025-08-06), using the EVcouplings package^23^ and a permissive domain/sequence bitscore threshold of 0.2. Hits were kept only if they covered the query at ≥60% sequence identity. SIFTS alignments mapped query residue indices onto each homolog PDB chain, and preferred biological assemblies were taken from PDBe^71^. For every mapped chain, distances were computed from the query-mapped residues to all other polymer and ligand entities in the assembly. Per query position, the reported contact is the minimum distance to a protein residue or ligand across homolog structures. Allowed ligands exclude a generic crystallography set (*e*.*g*., water molecules, ions, glycerol, PEG, EDO, DMSO). In some cases, selecting functional ligands required manual curation—for instance, vanadate ions can bind non-specifically but can occupy the active site in PTEN^59^. Regulatory domain interactions in Src were computed in an analogous way using structures representing different conformations.

### 4.5 A standardized scale to represent phenotypic contributions of variant effects

Given extensive prior work linking protein destabilization with low evolutionary selection^33,34^, and since structural stability is a *prerequisite* for function, we assume that the more deleterious a mutation is, the more likely it is that the consistency between predicted stability and evolutionary fitness explains its deleteriousness. We represent the overall mutational effect size using the predicted fitness from an evolutionary model transformed into a deleteriousness metric *w*, which is the min-max normalized fitness value across all deleterious mutants. Thus, *w* = 0 for the wild-type protein and *w* = 1 for the substitution with the minimum evolutionary fitness score.

We then combine scores from the deconvolution module (see Methods 4.3) with deleteriousness *w* using a weighting function dependent on the scale of the deconvolution scores. In other words, the predicted deconvolution effect is amplified with deleteriousness by defining the stability share *s*_*s*_ as

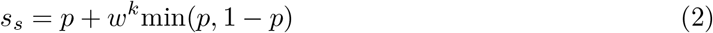

where *p* ∈ [0, 1] is the raw deconvolution score (see Fig. 2c), *w* ∈ [0, 1] is deleteriousness, and *k* ≥ 1 tunes the non-linearity of the amplification. The corresponding activity share *s*_*a*_ is simply the complement of the stability share:

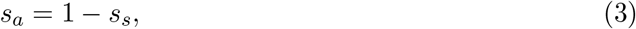

where *s*_*s*_ *∈* [0, 1] and *s*_*a*_ *∈* [0, 1].

This simple formulation seamlessly encodes the hierarchical effects of protein function. The correction term *w*^*k*^min(*p*, 1 − *p*) effectively amplifies the stability share of phenotypically consistent mutation effects proportionally to deleteriousness *w*. A greatly deleterious mutant (*w* → 1) that exhibits an inconsistency between stability and evolutionary model scores (*p* → 0) exerts its mutation effect almost exclusively by modulating activity (*s*_*a*_ ≈ 1, *s*_*s*_ ≈ 0)—a stable-but-inactive variant. In contrast, the effect of a perfectly consistent but deleterious mutant (*p* → 0.5, *w* → 1) is fully explained by stability (*s*_*a*_ ≈ 0, *s*_*s*_ ≈ 1), and a smooth gradient in *p* is obtained between these edge cases.

This convention also establishes a biologically informed gradient across mutational severity *w*, effectively mirroring our confidence in the deconvolution. As deleteriousness decreases, the stability and activity shares naturally converge toward their raw values, which are moderate at low deleteriousness regimes given the significant correlation of stability and activity for most targets (Fig. 2). In these cases, the detected inconsistencies between stability and evolutionary fitness dimensions indicate a marginal evidence for the phenotype that drives the mutation effect. This also reflects the added difficulty of separating the phenotypic contributions of cases of mild deleteriousness, and the potential for undeconvolvable mutation effects. The shape of this transition can be tuned with *k*, where higher values progressively restrict the weighting to the most catastrophic mutants.

### 4.6 Network routing module and phenotype enrichment test

We represented each protein as an undirected residue graph derived from EVcouplings probabilities^23^. EVcouplings fits a regularized pairwise Potts model to a MSA^42^ and summarises each pair by a scalar coupling score, which is then mapped to a probability of a structural contact^6^. These probabilities therefore capture coevolution between residues and are enriched for spatial contacts. In order to only keep pairwise relationships that could mediate biophysical mechanisms, couplings with a minimum heavy-atom distance above 5 Å were set to zero. To ensure that only high-confidence couplings are considered, pairs with probability *≥* 0.7 were kept as edges, and we assigned each edge a weight of 1 *− p*, where *p* is the EVcouplings probability^23^.

We denote by *information sinks* the subset of residues in substrate binding positions (minimum distance *<* 3.5 Å, see Methods 4.4), and we denote by *information sources* the subset of residues corresponding to the bottom 1% mutants by deconvolution score (see Methods 4.3). For every unique source-sink pair up to *k* = 5 lowest-cost simple paths were generated using Yen’s algorithm, thereby simultaneously minimizing edge number and cost^72^. Generating multiple paths allows sampling routes of alternative coupling strengths and lengths^73^. Given the high minimum coupling strength cutoff of 0.7, for each source-sink pair, resulting paths were ranked to select the path with the least edges. Only the top path for each activity modulator identified as an information source (allosteric ligand-binding site, protein-protein interface, or a site identified *de novo*) was tested. For NUDT15, a minimum edge count of 5 was imposed given the dense network of first- and second-degree source-sink pairs.

The enrichment for DMS phenotypes of path residues was evaluated through a Monte Carlo Randomization (MCR) test against random residue sets of the same length and comparable RSA. In this manner, the MCR test accounts for buriedness, which is a strong predictor of stability (Supp. Fig. 1). The empirical null distribution was built by drawing samples from a pool of residues with RSA values within a *±*0.1 window without replacement for each path constituent. Observed path residues were excluded from every null set. Residue-level phenotypes were then aggregated to a set statistic and compared to the null distribution. Significance values were computed as one-sided Monte Carlo with add-one continuity:

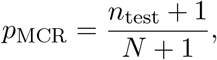

where *n*_test_ is the number of randomized null sets that achieve a set statistic equal to or more extreme than the observed set, and N is the total number of permutations (*N* = 1, 000). As a consequence, the maximum observable significance (*n*_test_ = 0) is *p*_MCR_ = 9.99 *×* 10^*−*4^.

Enrichment tests can be carried out for continuous DMS phenotypes or proportions of phenotypes of interest. For continuous stability, activity or model scores, the set score is the mean of the mean value over all single mutants evaluated at each site. For the binary case— wild-type-like stability or catastrophic mutants, where both activity and stability are below wild-type—the set score is the mean fraction of mutants with the phenotype of interest across positions in the set. In addition to *p*_MCR_ an empirical *Z*-score can be computed as:

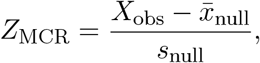

which represents the standardized observed effect size controlling for buriedness. The tests reported in the main text were repeated with two alternative nulls to assess the robustness of the associations. Both nulls were drawn from simple paths of the same length as the tested path on a structural contact graph. The first null comprised contact paths alone, and the second retained a path only when its mean RSA was within 0.1 of the mean for the tested path. Contacts were defined from the representative structures in Methods 4.1 as C*α*–C*α* pairs at a distance cutoff of *<* 8 Å. Where several structures were used, the distance was the minimum across structures. Results are shown in Supp. Table 3.

## 5 Data and code availability

Evolutionary models EVmutation2, EVE and ESM2 are available within the evedesign software suite^35^. EVcouplings and EVmutation are available both in evedesign and within its dedicated suite at https://evcouplings.org^23^. All steps of the pipeline can be accessed at https://github.com/sgb67/dualde, alongside examples and the outputs from the routing module used throughout the manuscript.

## 8 Supplementary Material

### 8.1 Supplementary Tables

**Supplementary Table 1:**
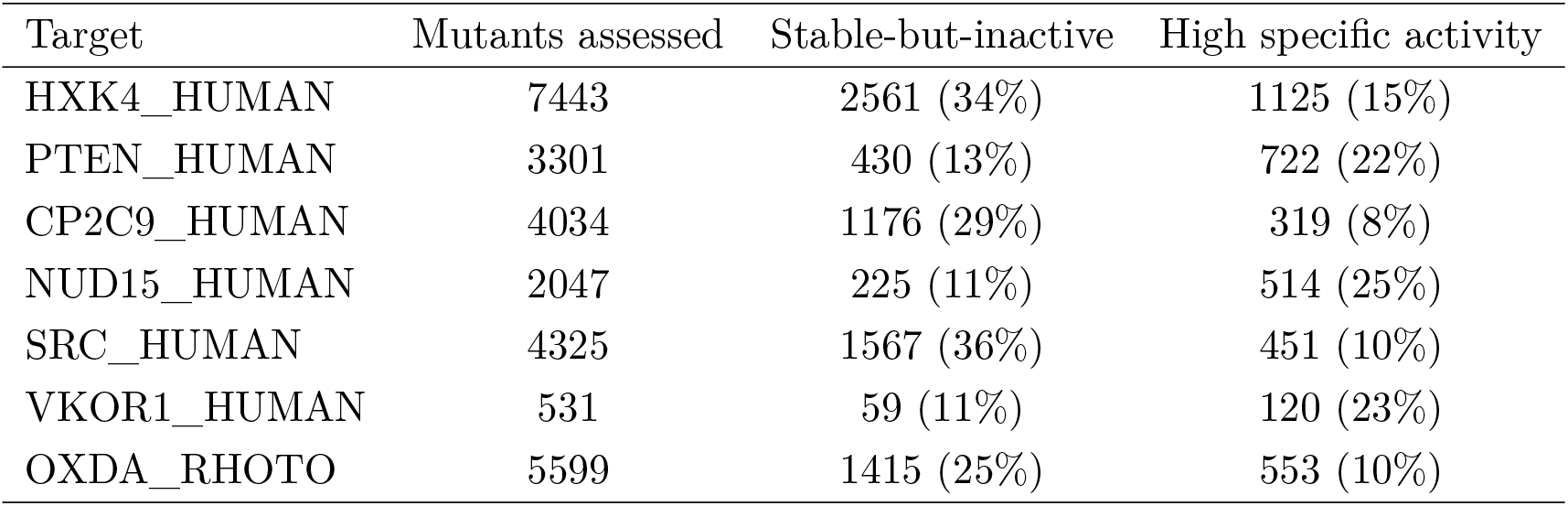
Number of mutants assessed for each target with the number of decoupled variants. Only positions that could be scored by evolutionary models and stability predictors were considered. For SRC, only the kinase domain was assessed^16^. Wild-type ranges were obtained from the original sources (see Methods 4.1).

**Supplementary Table 2:** All paths highlighted in the main text, including information sources and sinks, number of edges, and mean coupling strengths. All information sources were identified by the deconvolution module and were within the top 1% of deconvolved mutants. All potential information sinks were residues contacting the substrate or substrate analogs (*<* 3.5 Å). The path ranking reported is relative to all shortest paths drawn from that information source, across all possible sinks. For each unique pair, a maximum of *k* = 5 shortest-paths were generated, and paths were ranked by number of edges (see Methods 4.6). The total number of paths found for each information source is shown in parentheses. Since NUDT15 contained a dense network of high coupling strengths due to the proximity of the active site and dimerization interface, a minimum edge count of 5 was set for that target.

| Target | Modulator | Source | Sink | Edges | Mean coupling | Path rank |
| --- | --- | --- | --- | --- | --- | --- |
| GCK | Ligand | 451 | 169 | 7 | 0.82 | 1 (30) |
| GCK | Heterodimer | 198 | 168 | 12 | 0.94 | 1 (30) |
| OXDA | Homodimer | 185 | 335 | 8 | 0.98 | 1 (25) |
| NUDT15 | Homodimer | 82 | 15 | 6 | 0.99 | 1 (73) |
| SRC | Homodimer | 521 | 346 | 15 | 0.90 | 1 (10) |
| SRC | Regulatory domain | 368 | 346 | 14 | 0.84 | 1 (10) |
| PTEN | Catalytic switch | 77 | 126 | 5 | 0.82 | 1 (15) |

**Supplementary Table 3:** Phenotype enrichment test results for the residue sets from the top paths highlighted in the main text. The target, information source, phenotype to be tested for enrichment, and null distribution construction method are shown, together with the MCR p-value and empirical Z-scores for each test. RSA-matching corresponds to the test reported in the main text, where residues with a similar buriedness to each site in the selected top path are sampled to build the null distribution. The “random paths” test refers to a null distribution built with paths of the same length as the selected top path that are physically contiguous. The “paths + RSA” test refers to a null distribution built with paths with physically contiguous residues where the average RSA for each path is similar to that of the selected top path (see Methods 4.6). RSA similarity was set to *±*0.1 RSA. All nulls were built with the same seed. MCR, Monte Carlo Randomization.

| Target / Source | Phenotype | RSA matching |  | Random paths |  | Paths + avg. RSA |  |
| --- | --- | --- | --- | --- | --- | --- | --- |
|  |  | *p* | *Z* | *p* | *Z* | *p* | *Z* |
| GCK / Ligand | Stability | 0.007 | 2.436 | 0.031 | 1.720 | 0.054 | 1.666 |
| GCK / GKRP | Stability | 0.001 | 4.508 | 0.014 | 2.192 | 0.014 | 2.298 |
| OXDA / Dimer | Stability | 0.001 | 3.260 | 0.059 | 1.723 | 0.073 | 1.752 |
| NUDT15 / Dimer | Catastrophic | 0.047 | 1.662 | 0.080 | 1.566 | 0.125 | 1.149 |
| SRC / Various | Catastrophic | 0.002 | 2.692 | 0.059 | 1.840 | 0.099 | 1.344 |
| PTEN / Modulator | Stability | 0.005 | 2.997 | 0.094 | 1.325 | 0.033 | 1.833 |

**Supplementary Table 4:**
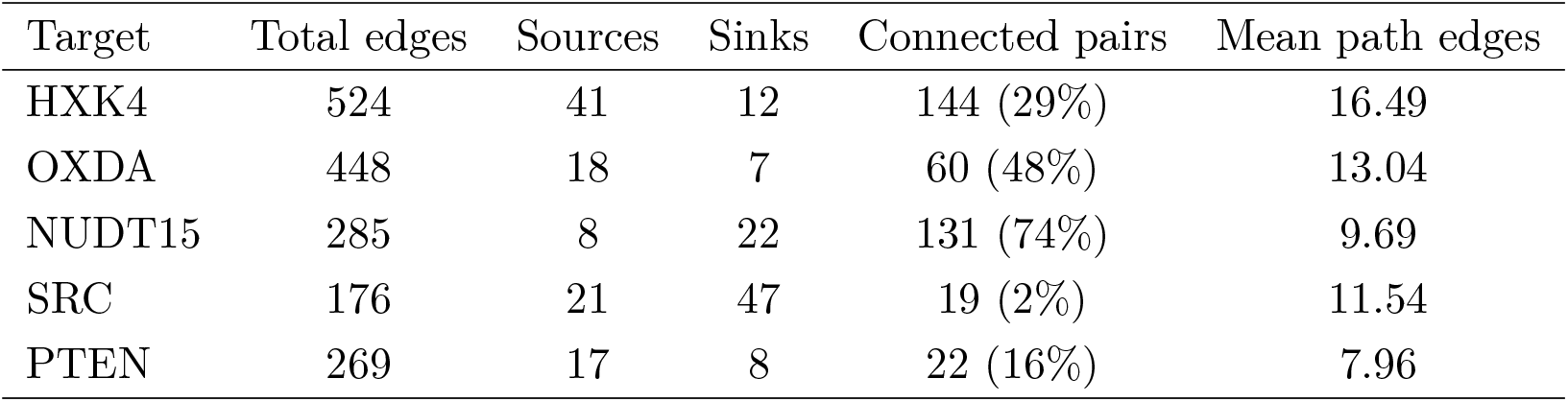
Coevolutionary information graph statistics from the routing module, including total number of edges, sources, sinks, connected pairs, and mean edge count amongst identified paths. Information sources were sites identified by the deconvolution module and were within the top 1% of deconvolved mutants. Information sinks were residues contacting the substrate or substrate analogs across homologs (*<* 3.5 Å, see Methods 4.4). The number of unique source-sink pairs connected by at least one allowed path is also shown, as well as the proportion it represents within all possible unique source-sink pairs in parentheses (see Methods 4.6). The variable proportion is expected due to the fact that the routing module results depend on a range of factors, including protein size, domain layout, alignment quality, and the location of the deconvolved mutant sites relative to the active site.

| Target | Total edges | Sources | Sinks | Connected pairs | Mean path edges |
| --- | --- | --- | --- | --- | --- |
| HXK4 | 524 | 41 | 12 | 144 (29%) | 16.49 |
| OXDA | 448 | 18 | 7 | 60 (48%) | 13.04 |
| NUDT15 | 285 | 8 | 22 | 131 (74%) | 9.69 |
| SRC | 176 | 21 | 47 | 19 (2%) | 11.54 |
| PTEN | 269 | 17 | 8 | 22 (16%) | 7.96 |

### 8.2 Supplementary Figures

**Supplementary Figure 1:**
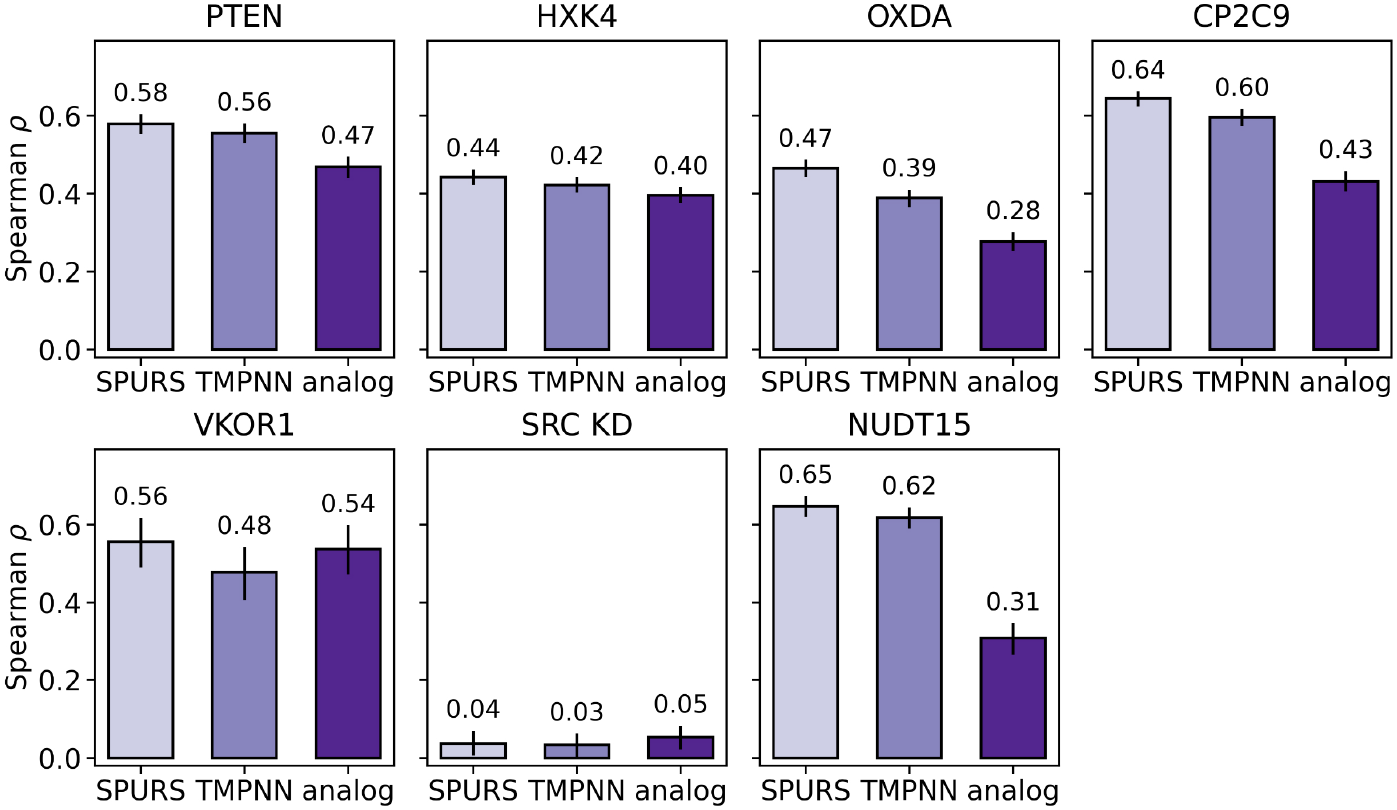
Spearman correlations of stability predictors with experimental stability by DMS for all enzymes in the validation set. SPURS^37^, ThermoMPNN^36^ (TMPNN) and the RSA analog (see Methods 4.1) are shown. 95% confidence intervals were obtained via non-parametric bootstrapping (*n* = 1, 000).

**Supplementary Figure 2:**
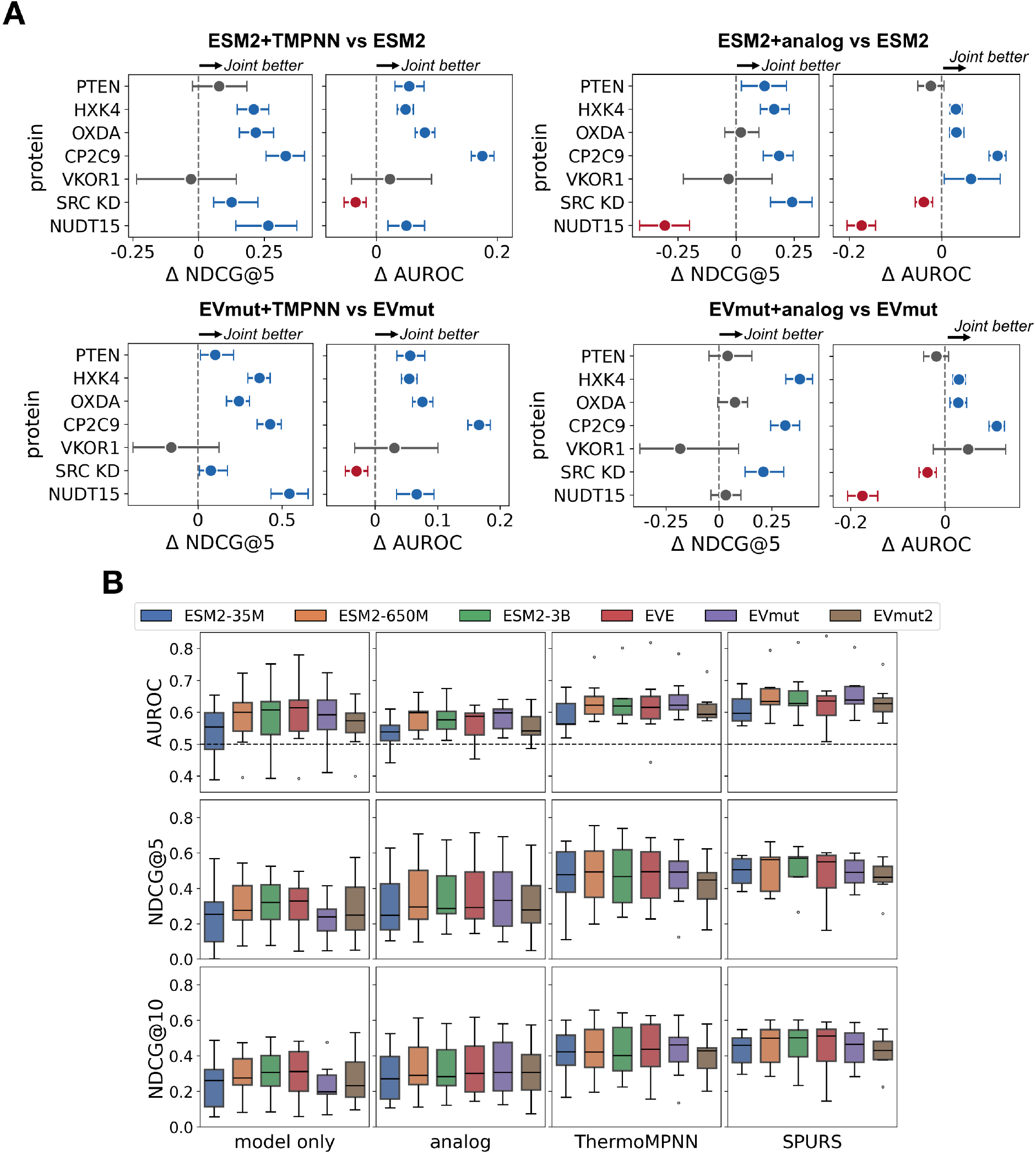
Recovery of stable-but-inactive variants across the validation set. (A) Head-to-head comparisons between joint models with ThermoMPNN (left) and the RSA analog (right) as the stability predictor and evolutionary models alone. The top row corresponds to ESM2 (650M) and the bottom to EVmutation. (B) AUROC, NDCG@5 and NDCG@10 distributions for all models, stratified by stability predictor and colored by evolutionary model choice. For SRC, only the kinase domain was processed. The difference in each metric is reported with 95% confidence intervals, which were obtained by non-parametric bootstrapping of 1,000 samples.

**Supplementary Figure 3:**
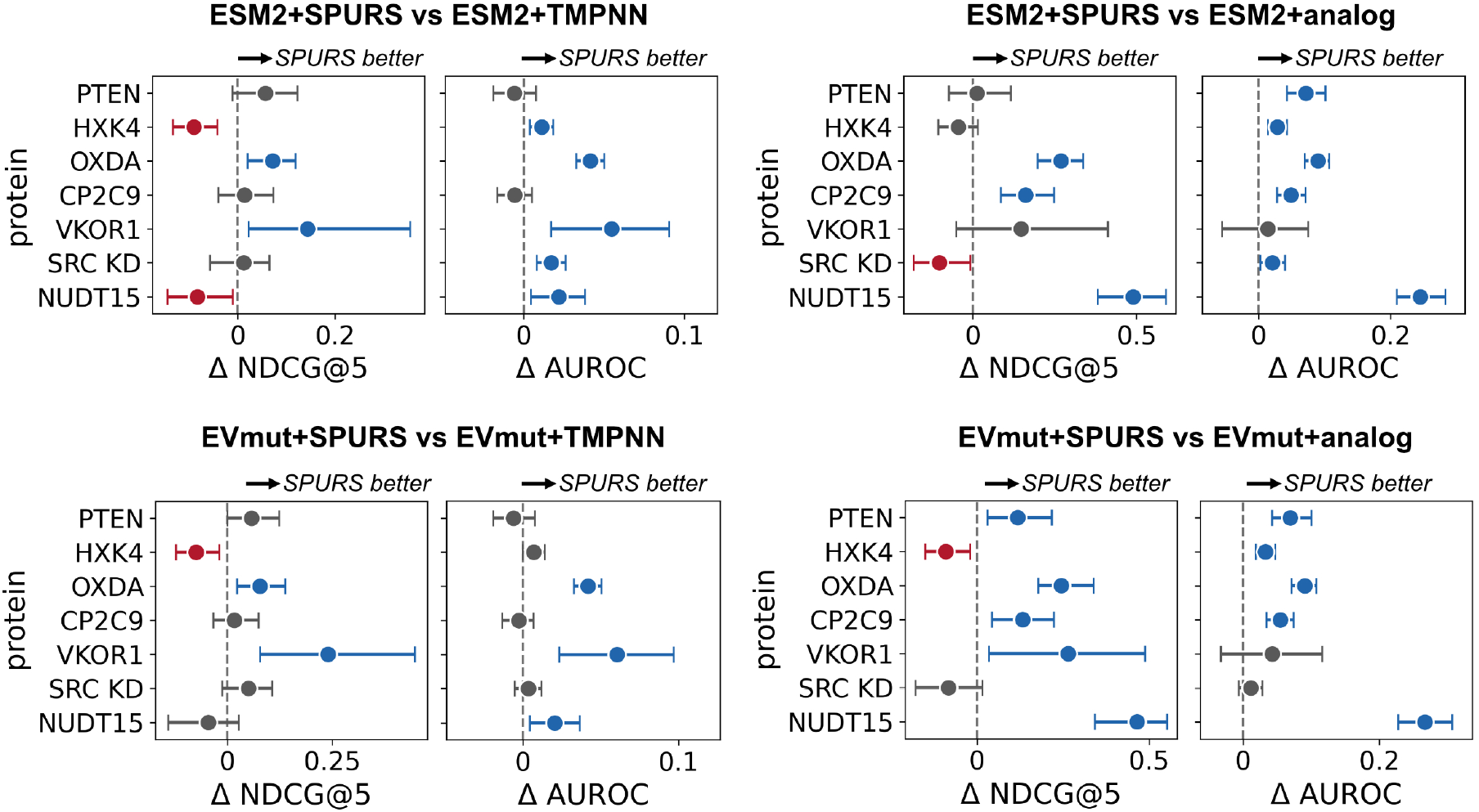
Head-to-head comparisons between joint models with SPURS against ThermoMPNN as the stability predictors (left) and SPURS against the RSA analog (right). The top row corresponds to ESM2 (650M) and the bottom to EVmutation. The difference in each metric is reported with 95% confidence intervals, which were obtained by non-parametric bootstrapping of 1,000 samples.

**Supplementary Figure 4:**
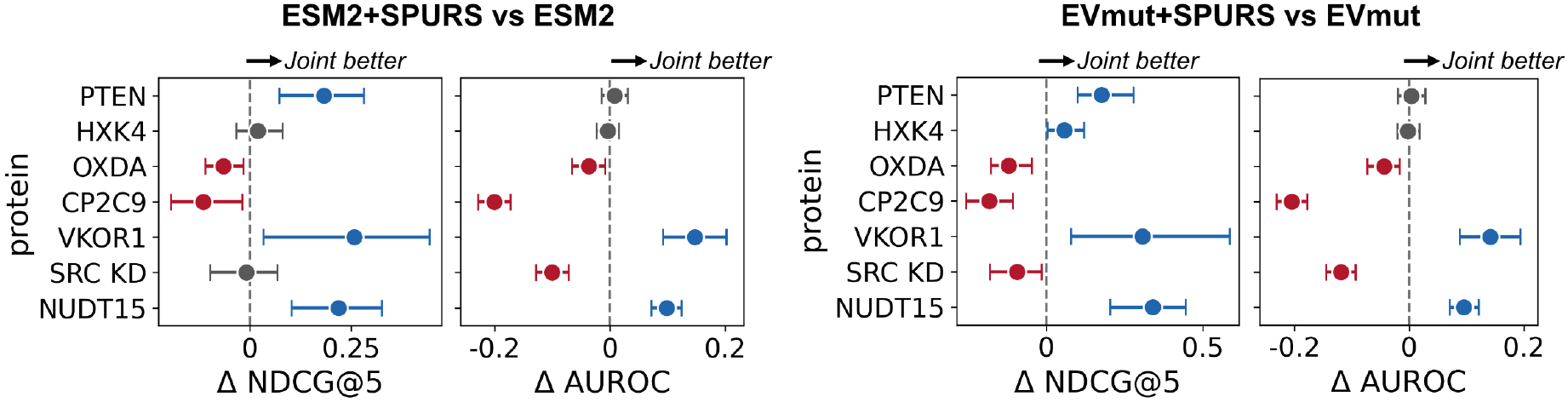
Head-to-head comparisons between joint models and evolutionary models alone for recovery of variants with high specific activity (see Methods 4.1). High model scores were screened to test for recovery. The difference in each metric is reported with 95% confidence intervals, which were obtained by non-parametric bootstrapping of 1,000 samples.

**Supplementary Figure 5:**
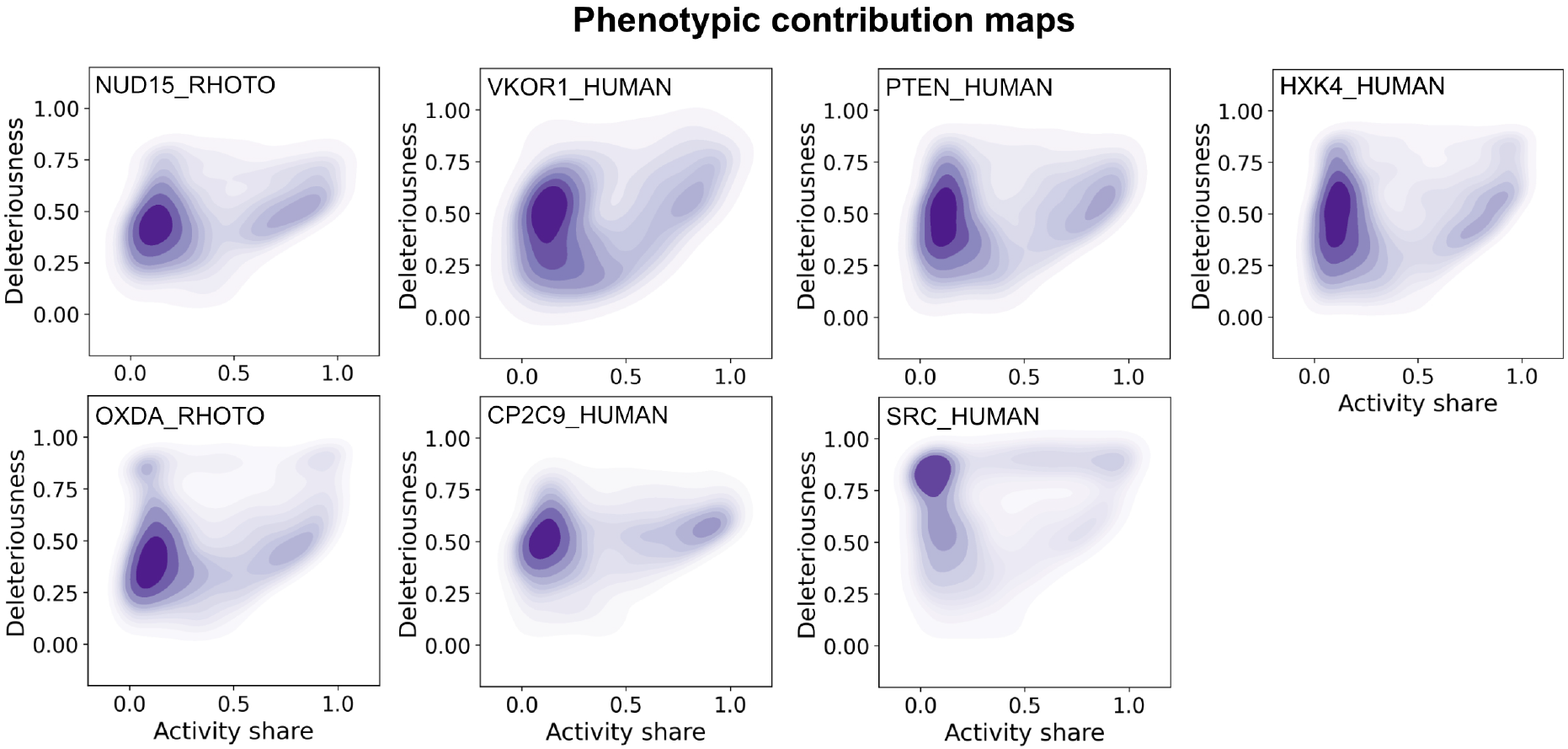
Phenotype deconvolution fingerprints for all enzymes in our validation set (see Methods 4.5). UniProt identifiers are shown within each plot. For SRC, only the kinase domain was processed.

**Supplementary Figure 6:**
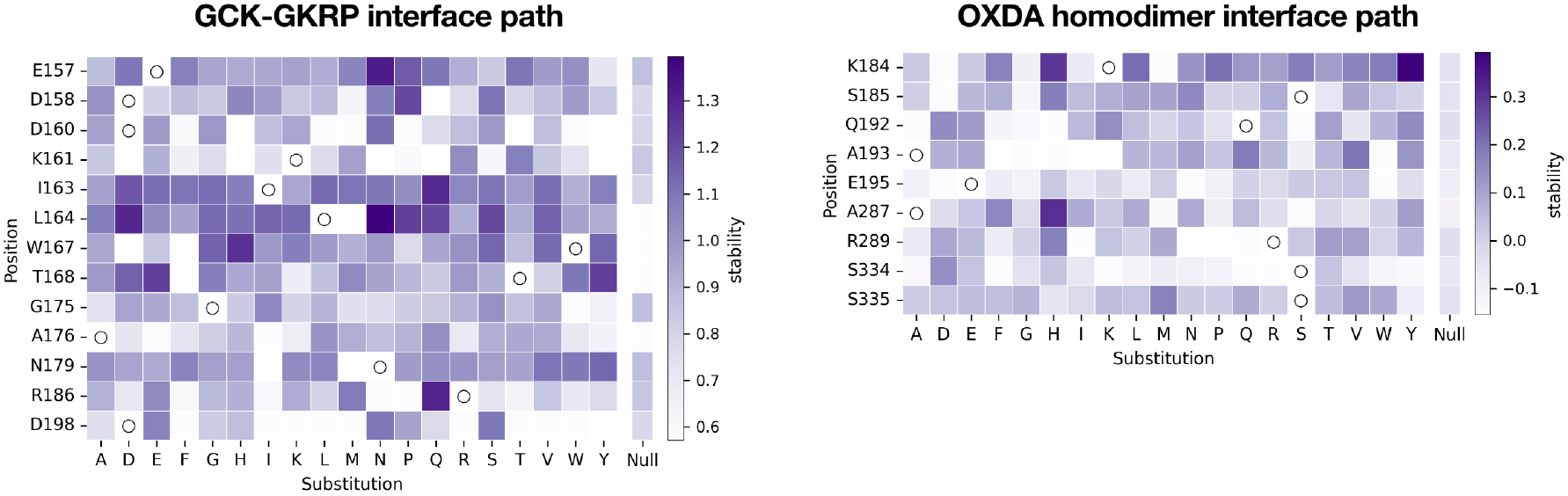
Stability profiles of path constituents for GCK-GKRP and OXDA dimeric interfaces compared to the null expected stability for a residue of similar buriedness (*±*0.1 RSA). RSA, Relative Solvent Accessibility.

